# Raman spectral analysis of microbial pigment compositions in vegetative cells and heterocysts of multicellular cyanobacterium

**DOI:** 10.1101/2023.02.13.527956

**Authors:** Jun-ichi Ishihara, Hiroki Takahashi

## Abstract

The one-dimensional multicellular cyanobacterium, *Anabaena* sp. PCC 7120, exhibits a simple topology consisting of two types of cells under the nitrogen-depleted conditions. Although the differentiated (heterocyst) and undifferentiated cells (vegetative cells) were distinguished by their cellular shapes, we found that their internal states, that is, microbial pigment compositions, were distinguished by using a Raman microscope. Almost of Raman bands of the cellular components were assigned to vibrations of the pigments; chlorophyll *a*, β-carotene, phycocyanin, and allophycocyanin. We found that the Raman spectral measurement can detect the decomposition of both phycocyanin and allophycocyanin, which are components of the light-harvesting phycobilisome complex in the photosystem II. We observed that the Raman bands of phycocyanin and allophycocyanin exhibited more remarkable decrease in the heterocysts when compared to those of chlorophyll *a* and β-carotene. This result indicated the prior decomposition of phycobilisome in the heterocysts. Moreover, the Raman bands of allophycocyanin were more decreased in heterocysts when compared to those of phycocyanin, suggesting that the decomposition of phycocyanin was more strongly suppressed than allophycocyanin in heterocysts. We show that the Raman measurement is useful to detect the change of pigment composition in the cell differentiation.

## Introduction

Cyanobacterium is a gram-negative prokaryote that carries out oxygenic photosynthesis [1]. Among of them, *Anabaena* sp. PCC 7120 (hereafter named *Anabaena*) is a filamentous and multicellular cyanobacterium [2–4] (Figure 1A). The filament is composed of a lot of cells connected in a one-dimensional manner.

**Figure 1.**
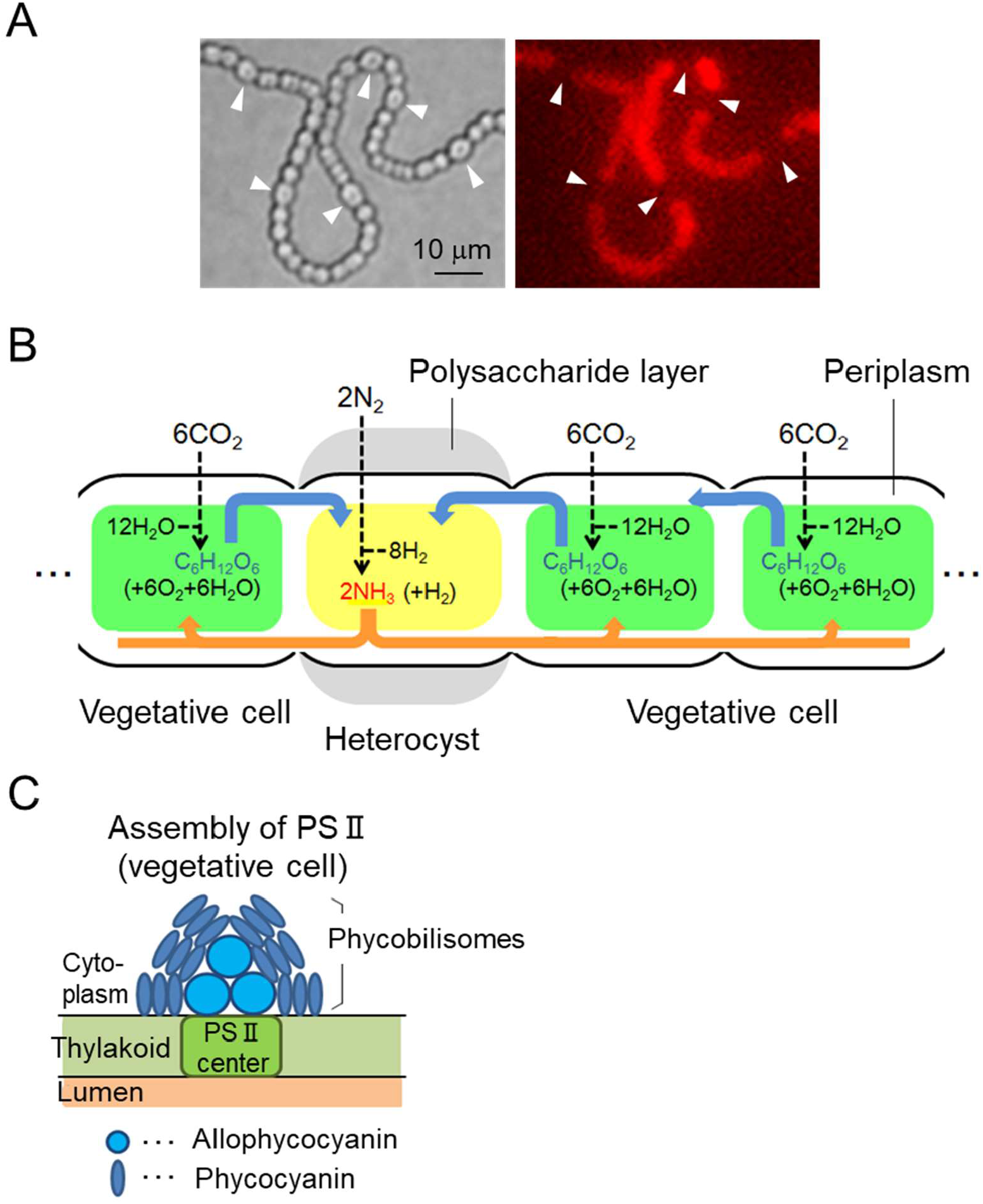
Schematic figure of the heterocyst pattern formation of *Anabaena* sp. PCC 7120 (*Anabaena*) under nitrogen-depleted conditions. (A) Photomicrographs of the heterocyst pattern. Heterocysts are observed as expanded cells with degraded phycobilisome complex. Left and right panels show a bright field and a phycobilisome fluorescence micrograph, respectively. White arrows indicate heterocysts. Scale bar is 10 μm. (B) Schematic representation of metabolic transfer between a heterocyst and neighboring vegetative cells. While heterocysts produce nitrogen compounds by the nitrogen fixation, vegetative cells produce carbohydrates by the photosynthesis. The heterocyst and neighboring vegetative cells exchange the carbohydrates and nitrogen compounds through periplasm surrounding the cells. (C) Schematic representation of phycobilisome (a photosystem II light harvesting complex). Six pieces of phycocyanin aggregates locate outside, surrounding a single piece of allophycocyanin aggregate for a single unit of the phycobilisome.

In nitrogen-depleted culture medium, *Anabaena* initiates cellular differentiation [2–8]. The differentiation occurs at an interval of approximately ten cells along the filament. Differentiated and undifferentiated cells are named as heterocysts and vegetative cells, respectively. A heterocyst is easily distinguished by using a standard microscope because it is larger in size and rounder in shape. Once a cell is fully differentiated, it never reverts vegetative cell. As the number of vegetative cells increases by the cell division, a new heterocyst is differentiated approximately midway between two older heterocysts. A heterocyst fixes dissolved nitrogen to the reduced nitrogen species, e.g., ammonium ion [6, 7]. In addition, the light-harvesting phycobilisome complex is chemically decomposed or inactivated, leading to suppress the oxygen-evolving photosystem II (PSII) [6–10]. This is because a nitrogen-fixing enzyme, nitrogenase, is sensitive to oxygen. On the other hands, vegetative cells are dedicated for the photosynthesis and contain both PSI and PSII, while heterocysts mainly contain PSI. As the photosynthesis and nitrogen fixation are incompatible in the same cell, the heterocysts and vegetative cells exchange metabolites produced by the nitrogen fixation or photosynthesis with the neighboring cells [4–7] (Figure 1B).

The remarkable differences of the photosynthetic systems between vegetative cells and heterocysts have been studied by using a fluorescence microscope [9–13]. However, observations and analysis on the heterocyst-forming cyanobacteria by using a Raman microscope have been limited [14–16]. Recently it is possible to measure Raman spectra of a single live cell [17–18]. Biomolecules such as proteins, nucleic acids, lipids, etc., that is, atoms combined with electrons, vibrate with different wavenumbers. These oscillators interact with the light and are detected as Raman scattering.

In this study, we analyzed the distribution of biomolecules including small molecules in living cells in a non-invasive and non-labeling manner using Raman signals. We found that vegetative cells and heterocysts were classified by 16 Raman bands in the Raman spectra by excitation at 785 nm laser. These Raman bands were assigned to vibrational modes of resonance Raman bands of four pigments known as light-harvesting pigments, chlorophyll *a*, β-carotene, phycocyanin, and allophycocyanin. We also found that the components in the phycobilisome in PSII, that is, phycocyanin and allophycocyanin, were detected by analyzing the high resolution of Raman bands with sharp band widths. We calculated the correlations of band intensities among four pigments in vegetative cells and heterocysts. As a result, the intensities of Raman bands of phycocyanin and allophycocyanin in the heterocysts were remarkably decreased when compared to those of chlorophyll *a* and β-carotene. Our result shows good correspondence with the previous studies that phycobilisome in PSII is decomposed during the differentiation process [9, 10]. Moreover, the Raman bands of allophycocyanin exhibited more remarkable decrease in heterocysts when compared to those of phycocyanin. This result suggests that the decomposition of phycocyanin was suppressed in heterocysts. A Raman microscope helps us to analyze the change of the PSII components before and after the differentiation.

## Materials and Methods

### Bacterial strains and culture

*Anabaena* sp. PCC 7120 (wild type) were grown in 25 ml of BG-11_0_ (lacking sodium nitrate) liquid medium in 50 ml flasks at 30 °C under illumination with white fluorescent lamps (FL30SW-B, Hitachi co.) at 45 μM photons m^-2^s^-1^. The culture was being shook at 120 rpm until an optimal density at 730 nm (OD_730_) reached about 0.4–0.5. The liquid culture was washed three times with BG11_0_, diluted to an OD730 of ∼0.2, and underlain beneath a fresh BG-11_0_ solid medium plate containing 1.5% agar solution (Becton, Dickinson and company, USA) with a bottom dish glass. The sample was placed in a Raman microscope (as mentioned below) kept at 30 °C under illumination with white fluorescent lamps at 45 μM photons m^-2^s^-1^.

### Reference pigments

Raman spectra of pigments were obtained by using chlorophyll *a* (Sigma-Aldrich, C6144-1MG), β-carotene (Wako Pure Chemical Industries, 035-05531), phycocyanin (Sigma-Aldrich, P2172), and allophycocyanin (Sigma-Aldrich, A7472). To measure the spectra, these pigments were dissolved in BG11_0_ liquid medium and underlain beneath a fresh BG-11_0_ solid medium plate containing 1.5% agar solution with a bottom dish glass. The condition of measurement was the same as that mentioned above.

### Raman microscope and spectral pre-treatments

In Via confocal Raman spectrometer equipped with a CCD camera (inVia Reflex, Renishaw co.) was used to measure the Raman spectrum. The excitation wavelength was at 785 nm. We measured Raman spectra of individual cells by selecting the central points of the cells. A typical Raman spectrum of a small confocal volume in the cytoplasm (horizontal diameter, ∼ 1 μm) of a single living vegetative cell (∼ 3 μm diameter) yields a sufficient signal-to-noise ratio for analysis (∼1 s per pixel, with a 785 nm laser at ∼20 mW directed at the confocal volume). In this study, the baselines of Raman spectra were corrected. The baseline-corrected Raman spectrum y’(v) was calculated as *y*’(*v*) = *y*(*v*) – *y*_poly_(*v*), in which *y*_poly_(*v*) is a fitted polynomial curve constructed with the following procedures. (i) For a spectrum truncated between the minimum Raman shift position *v*_min_ and the maximum position *v*_max_, the degree of the function *d* was selected to fit the baseline using a polynomial function (this time *d*=3). (ii) Using the least squares method, the polynomial function *y*_poly_ was first fitted to the Raman spectrum *y*. (iii) The Raman spectrum *y* was divided into upper and lower parts, relative to the fitted baseline *y*_poly_. (iv) The number of data points on the upper side of y was designated *N*_A_, and the number on the lower side of *y* was designated *N*_B_. If *N*_A_ < *N*_B_, the upper part of *y* was removed from the whole of *y*, and the Raman spectrum y was replaced with the lower part of the spectrum. Then, procedure (ii) was repeated. When *N*_A_ ≥ *N*_B_, the baseline was considered the best fit and optimal.

## Results

### Raman Spectral Measurements of Vegetative Cells, Heterocysts, and Four Pigments

The average Raman spectra of vegetative cells and heterocysts are shown in Figures 2A and B. The procedure to obtain the Raman spectrum is explained in Method section. Here briefly, fifteen vegetative cells (or fifteen heterocysts) were selected from three *Anabaena* filaments, and the fifteen Raman spectra were measured for every single cell. The intensity values of a Raman spectrum in the region from 990 to 1770 cm^−1^ were normalized to unity. The fifteen normalized spectra of the vegetative cells (or heterocysts) were averaged shown in Figures 2A and B.

**Figure 2.**
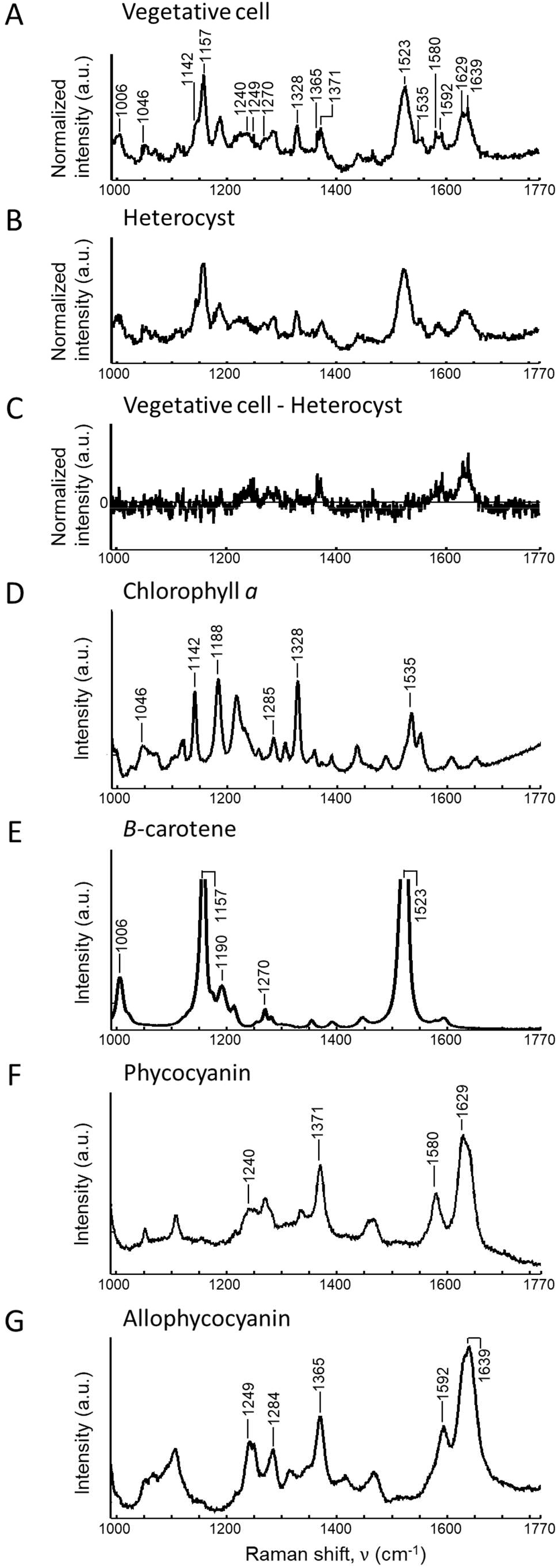
Raman spectra of living cells and photosynthetic pigments obtained with excitation at 785 nm. (A, B) The normalized intensities of Raman spectra of the vegetative cells and heterocysts. To calculate the Raman spectra, fifteen vegetative cells and fifteen heterocysts were selected from three *Anabaena* filaments, respectively. The Raman bands labeled by arrows in (A) were representative bands assigned to vibrations of chlorophyll *a*, β-carotene, phycocyanin, and allophycocyanin. These bands were also selected in Figure 3. (C) The difference between the Raman spectra (A) and (B). The Raman spectrum in Figure 2B was subtracted from that in Figure 2A. (D–G) The Raman spectra of chlorophyll *a*, β-carotene, phycocyanin, and allophycocyanin, respectively. The labeled Raman bands were the resonance bands assigned to vibrational modes from the previous studies [19–31].

The number of Raman band peaks in vegetative cells was almost identical to that in heterocysts (it was 16 bands), and the band positions in vegetative cells were nearly the same as those in heterocysts (the difference was within 2 cm^−1^) (Figures 2A and B). We found that the intensities of several Raman bands in vegetative cells were different from those in heterocysts. To show the difference of spectral features more clearly, we conducted that the Raman spectrum in heterocysts was subtracted from that in vegetative cells. As a result, the intensities of several peaks were different, i.e., 1240, 1249, 1365, 1371, 1580, 1592, 1629, and 1639 cm^−1^ (Figure 2C). This indicates that these peaks are potential differentiation markers for *Anabaena*. That is, heterocysts can be distinguished from vegetative cells solely using a Raman microscope.

Next, we measured four pigments in *Anabaena* by Raman microscope, that is, chlorophyll *a*, β-carotene, phycocyanin, and allophycocyanin (Figures 2D–F). By comparing the spectra with those in vegetative cells and heterocysts, almost all of the observed Raman bands of the vegetative cells and hetereocysts were explained (Figures 2A and B). When a living cell was exposed by the 785 nm laser light, spontaneous Raman bands derived from biomolecules such as lipids, proteins, and nucleic acids were assumed to be detected. In *Anabaena*, these biomolecules were not observed; instead, Raman bands of the chlorophyll *a*, β-carotene, phycocyanin, and allophycocyanin were detected. We consider that the Raman signals of Raman bands of four pigments were amplified due to resonance Raman effect. According to vibrational modes of resonance Raman bands of four pigments [19–31], we assigned the observed band peaks (Figures 2D–G). The Raman bands of four pigments were as follows: 1046, 1142, 1188, 1285, 1328, and 1535 cm^−1^ for chlorophyll *a*, 1006, 1157, 1190, 1270, and 1523 cm^−1^ for β-carotene, 1240, 1371, 1580, and 1629 cm^−1^ for phycocyanin, and 1249, 1284, 1365, 1592, and 1639 cm^−1^ for allophycocyanin (Figures 2D–G).

### Comparison of the normalized band intensities between vegetative cells and heterocysts

To elucidate the difference of composition of four pigments between vegetative cells and heterocysts, we compared the normalized intensities of the Raman bands of vegetative cells with those of heterocysts (Figure 3). The band positions in the Raman spectra of vegetative cells and heterocysts were almost identical to those in the Raman spectra of four pigments. Some Raman bands in the Raman spectra of the living cells were assigned to more than two vibrational modes of resonance Raman bands of pigments. For example, the Raman band at 1188 cm^−1^ in the Raman spectrum of chlorophyll *a* was very closed to that at 1190 cm^−1^ in the spectrum of β-carotene. Moreover, the Raman band at 1285 cm^−1^ in the Raman spectrum of chlorophyll *a* was also very closed to that at 1284 cm^−1^ in the spectrum of allophycocyanin. Consequently, the sixteen major bands (four times four) were selected for chlorophyll *a*, β-carotene, phycocyanin, and allophycocyanin.

**Figure 3.**
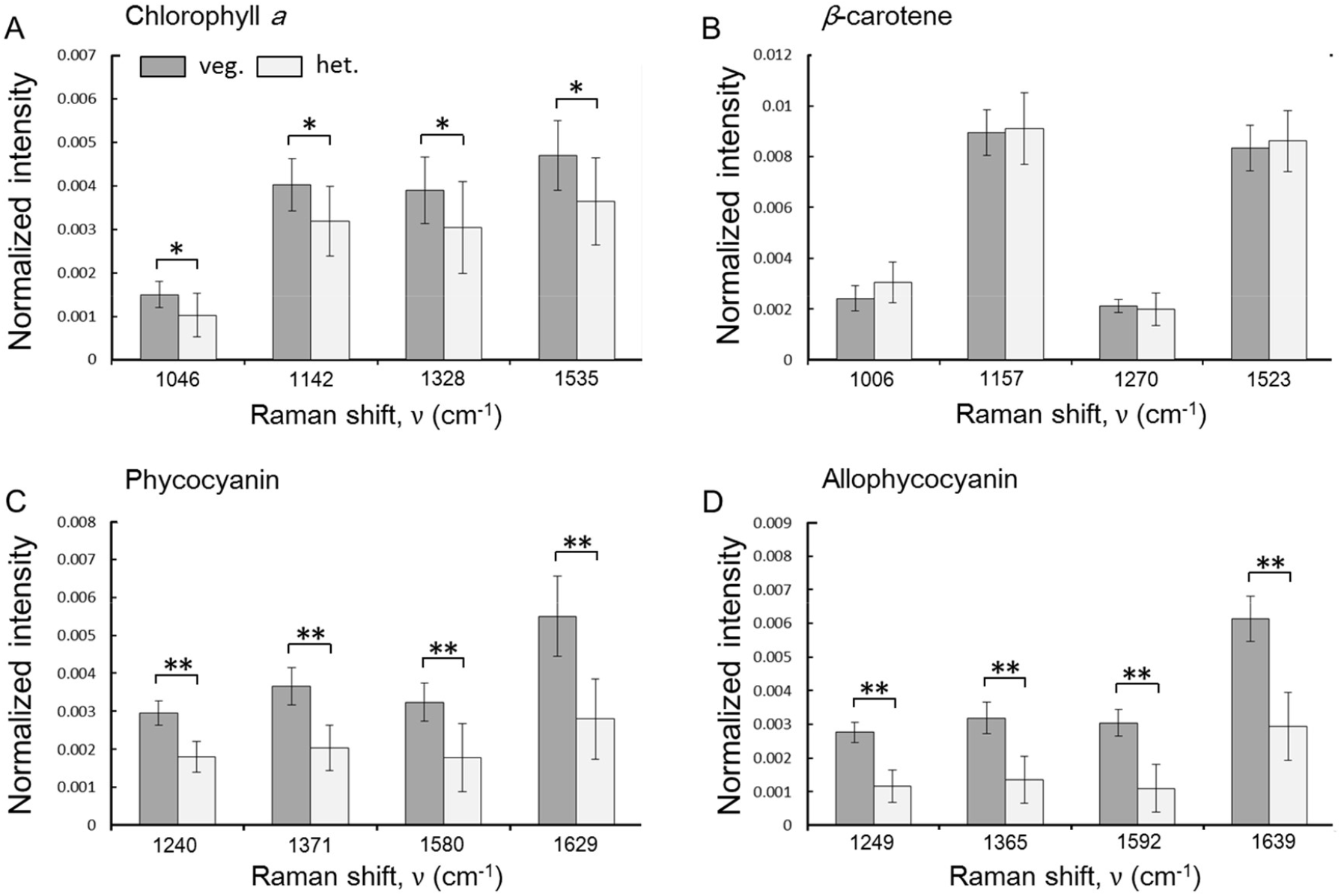
Comparisons of the normalized band intensities between the vegetative cells and the heterocysts, prepared based on the spectra in Figure 2A and B. The sixteen major bands (four times four) were selected for the chlorophyll *a*, β-carotene, phycocyanin, and allopycocyanin, respectively. Gray and black graphs show the normalized band intensities of the vegetative cells and heterocysts at each Raman shift (cm^−1^). Error bar indicates the standard deviations (*n*=15). Statistical difference is indicated by the following symbols. *: *p*<0.01, **: *p*<0.005.

All the band intensities of chlorophyll *a* of hetrocysts were decreased comparing with those of the vegetative cells (*p*<0.01, Welch’s *t*-test) (Figure 3A). Similarly, all the band intensities of phycocyanin and allophycocyanin of hetrocysts were decreased comparing with those of vegetative cells (*p*<0.005) (Figures 3C and D). However, the Raman band intensities of β-carotene were almost identical between the vegetative cells and heterocysts (Figure 3B), suggesting that β-carotene is chemically stable. It is known that while the amount of β-carotene is constant through the differentiation [9, 10, 16], the chlorophyll *a*, phycocyanin, and allophycocyanin are chemically decomposed, leading to impaired PSII. Therefore, we considered that the differences of the Raman bands of the chlorophyll *a*, phycocyanin, and allophycocyanin between the vegetive cells and heterocysts were caused by the chemical decompositions (a chemical decomposition reduces the amount of the target molecule and the corresponding Raman intensity).

Here, we found that the Raman band intensities of phycocyanin and allophycocyanin were largely decreased after the differentiation when compared to those of chlorophyll *a* (Figures 3A, C, and D). We considered that this is because chlorophyll *a* is a component of both the PSI and PSII, but phycocyanin and allophycocyanin, which form the phycobilisome as a protein complex unit, exist mainly in the PSII. Therefore, the selective decomposition of the PSII in the heterocyst resulted in the remarkable decrease of Raman bands of phycocyanin and allophycocyanin. However, the Raman band intensities of phycocyanin and allophycocyanin were far from zero in the heterocysts, suggesting that phycobilisome existed in the heterocysts. The gap of the values was explained in the Discussion section by referring latest studies.

Here, we focused on the normalized Raman band intensities of phycocyanin and allophycocyanin in the vegetative cells and heterocysts. After the differentiation, the four band intensities of phycocyanin were decreased by 38.2, 44.2, 44.3, and 48.2% at 1240, 1371, 1580, and 1629 cm^−1^, respectively. On the other hands, the four band intensities of allophycocyanin were decreased by 58.2, 57.7, 64.1, and 52.1% at 1249, 1365, 1592, and 1639 cm^−1^, respectively. That is, allophycocyanin was more decomposed in the heterocysts. Watanabe et al. reported that allophycocyanin was decomposed in preference to phycocyanin because a unique type of phycobilisome including phycocyanin was isolated from heterocysts of *Anabena* sp. PCC 7120 [32]. Watanabe et al. explains such new type of antenna complex may play an important role on light harvesting in PSI-driven cyclic electron transport to facilitate nitrogen-fixation and other reactions [32].

### Pigment Compositions and Correlations in the same cell

Correlations among intracellular pigment compositions were analyzed by selecting a representative Raman band among the four major bands (Figure 3) in each pigment. For example, in the case of chlorophyll *a*, the normalized band intensity at 1328 cm^−1^ was highly correlated to the other three normalized intensities at 1046, 1142 and 1535 cm^−1^ among 15 vegetative cells (*r*=0.71∼0.72, Supplemental Figure 1). These results suggest that the quantitative variation of chlorophyll *a* was reflected as the normalized band intensities in the living cells. Hereafter, the normalized band intensity at 1328 cm^−1^ was selectively used as the signal of chlorophyll *a*. In the case of β-carotene, phycocyanin, and allopycocyanin, the normalized band intensities at 1523, 1629, and 1639 cm^−1^ were also highly correlated to the other three normalized band intensities in each pigment (*r*=0.71∼0.74, Supplemental Figure 1). Thus, the normalized band intensities at 1523, 1629, and 1639 cm^−1^ were used as the representative signals of β-carotene, phycocyanin, and allophycocyanin, respectively.

In vegetative cells, almost no correlations (*r*=−0.19∼0.03) were found between β-carotene and other three pigments (Figures 4A, D, and E). That is, the amount of β-carotene was unrelated to that of other pigments in the PSI and PSII. On the other hands, moderate correlations (*r*=0.42 and 0.63) were found between chlorophyll *a* and the other two pigments in the phycobilisome (phycocyanin and allophycocyanin), respectively (Figures 4B and C). We consider that this correlation was observed because chlorophyll *a*, phycocyanin, and allophycocyanin are the components of PSII. Especially, high correlation (*r*=0.70) was found between phycocyanin and allophycocyanin (Figure 4F). This is probably related to the fact that six pieces of phycocyanin aggregates locate outside, surrounding a single piece of allophycocyanin aggregate for a single unit of the phycobilisome (Figure 1C). That is, the quantitative ratio of phycocyanin to allophycocyanin was tightly regulated in the phycobilisome.

**Figure 4.**
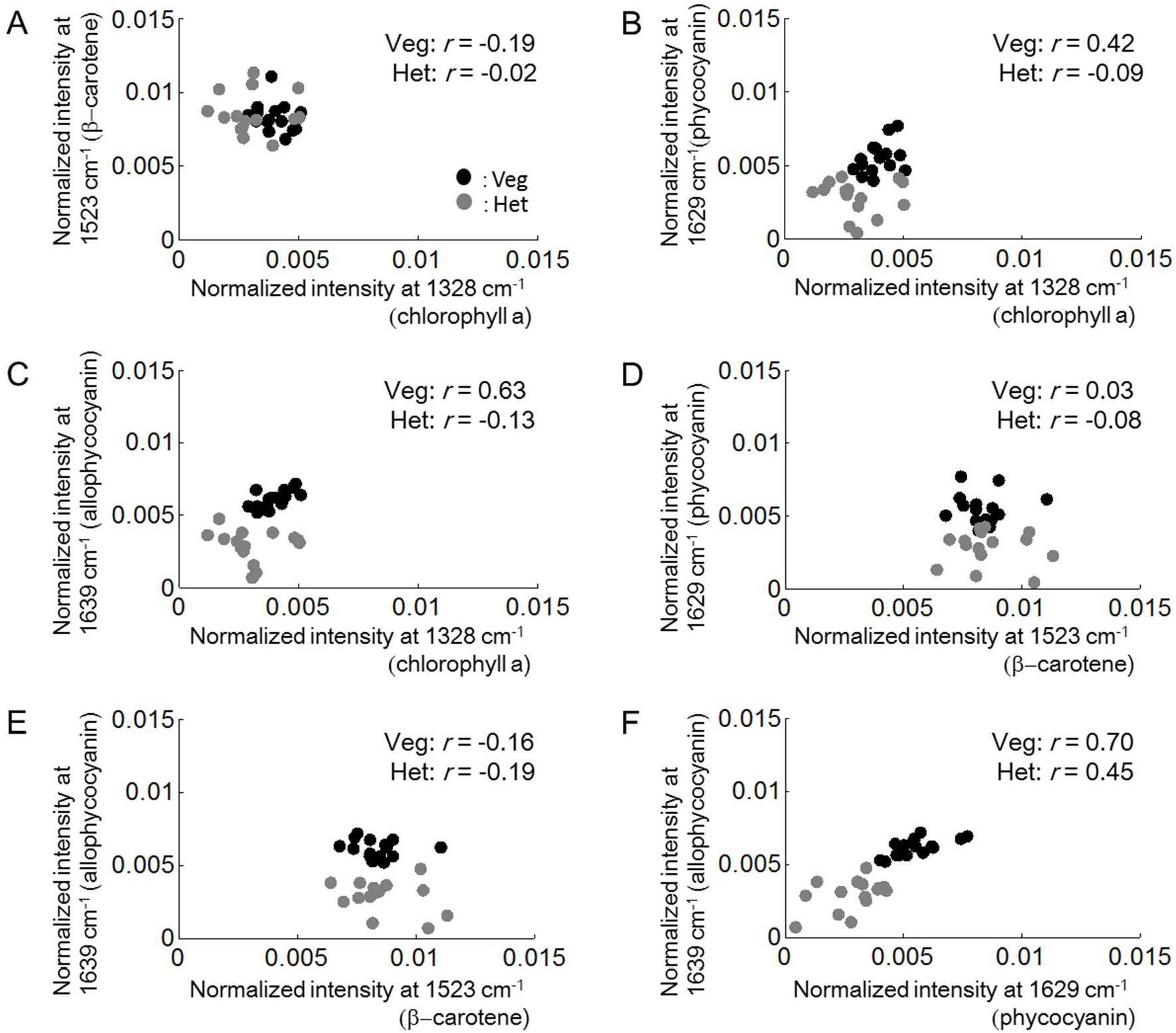
Correlation plots of the normalized band intensities between the pigments in the same vegetative cells and heterocysts. All plotted normalized band intensities were calculated by using the Raman spectra of the 15 vegetative cells and 15 heterocysts, which were also used in Figure 2 and Figure 3. Black and gray plots represent the band intensities of the vegetative cells and heterocysts. Numbers in all panels are correlation efficient among vegetative cells or heterocysts.

In heterocysts, we observed insignificant correlations (*r*=−0.09 and −0.13) between chlorophyll *a* and other two pigments in the phycobilisome (phycocyanin and allophycocyanin), respectively (Figures 4B and C). Moreover, the correlation between phycocyanin and allophycocyanin was moderate (*r*=0.45) when compared to the correlation value in the vegetative cells (Figure 4F). We considered that the PSII is decomposed in the heterocyst, and therefore, the correlations observed in the vegetative cells were disappeared or much weaker.

## Discussion

We measured the Raman spectra of the vegetative cells and heterocysts in *Anabaena* filaments. The Raman bands in the spectra were assigned to vibrations of chlorophyll *a*, β-carotene, phycocyanin, and allophycocyanin. In the heterocysts, the Raman band intensities of chlorophyll *a*, phycocyanin, and allophycocyanin were significantly decreased (Figures 3A, C, and D). We considered that this is because chlorophyll *a*, phycocyanin, and allophycocyanin are chemically decomposed with the PSII unfunctionalized through the differentiation. Especially, in the heterocysts, the band intensities of phycocyanin and allophycocyanin were remarkably decreased when compared to those of chlorophyll *a*. That is, we can distinguish the vegetative cells and heterocysts by analyzing the band intensities of phycocyanin and allophycocyanin. In the heterocysts, it is known that phycobilisome in PSII was decomposed [9, 10]. Therefore, we considered that the decrease of the band intensities of phycocyanin and allophycocyanin was due to the decomposition of phycobilisome in the heterocysts.

However, the band intensities of phycocyanin and allophycocyanin were not zero in the heterocysts, suggesting that phycobilisome was not perfectly decomposed. In heterocysts, efficient energy transfer has been proposed to occur from phycocyanin to PSI to facilitate nitrogen fixation and other reactions, but the transfer mechanism and structural details had been unknown [33]. Watanabe et al. reported an isolation of a unique phycobilisome-PSI supercomplex from heterocysts of *Anabaena* sp. PCC 7120 [32]. Biochemical and spectral analysis revealed that phycocyanin was included mainly in this type of phycobilisome, which was functionally connected to the PSI tetramer via a new connecting component, CpcL [32]. Therefore, phycocyanin was not decomposed perfectly and continued to exist in heterocysts. Watanabe et al. also reported that allophycocyanin was decomposed in preference to phycocyanin [32]. Our analysis corresponded to this result well because the Raman bands of allophycocyanin showed more remarkable decrease after the differentiation when compared to those of phycocyanin. This result suggests that phycocyanin and allophycocyanin were decomposed or restored to PSI by different mechanisms, respectively. As a future work, it should be addressed how phycocyanin and allophycocyanin are liberated from PSII, and how they are decomposed or restored in the heterocysts.

In this study, we distinguished the differentiated cells (heterocysts) from the undifferentiated cells (vegetative cells) by analyzing the pigment composition from the Raman spectra. The measurement of the Raman spectrum shown in this study is a useful technique to analyze the intracellular chemical composition without external probes. This methodology could have the potential to apply to other purpose, for example, the selection of focal bacterial cells from environments.

## Acknowledgements

We thank Dr. Shin-ichi Morita (Tohoku University), Dr. Sota Takanezawa (Nikon), Dr. Yasushi Sako (RIKEN), and Dr. Hideo Iwasaki (Waseda University) for valuable comments and advice. This work was supported by MEXT KAKENHI (11J06592, 17K15151, 22K15152) to J.I.

## Author contributions

JI conceived the project, performed the experiments, and analyzed the data. JI and HT wrote the manuscript and approved the submitted version.

## Figure and Supplementary Figure legends

**Figure S1.**
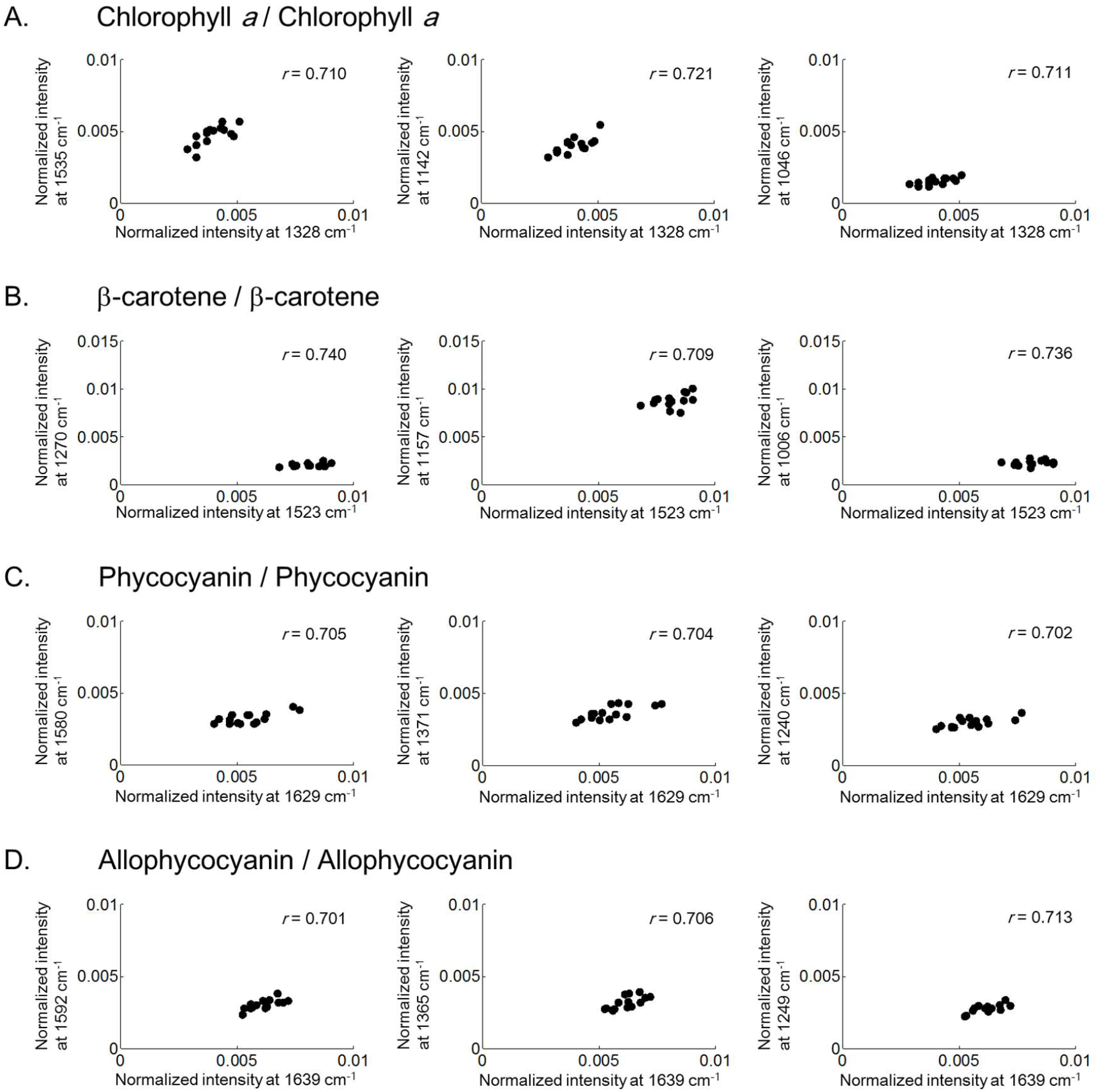
Correlation plots of the normalized band intensities assigned to the same pigments in the same vegetative cells. All plotted normalized band intensities were calculated by using the Raman spectra of the 15 vegetative cells, which were also used in Figure 2 and Figure 3. Numbers in all panels are correlation efficient among vegetative cells.

## References

1. Kumar, K., Mella-Herrera, R.A., and Golden, J.W. (2009). Cyanobacterial Heterocysts. Cold Spring Harbor Perspectives in Biology 2, a000300–314.

2. Wilcox, M., Mitchison, G.J., and Smith, R.J. (1973a). Pattern formation in the blue-green alga Anabaena. I. Basic mechanisms. J. Cell Sci. 12, 707–723.

3. Wilcox, M., Mitchison, G.J., and Smith, R.J. (1973b). Pattern formation in the blue-green alga Anabaena. II. Controlled proheterocyst regression. J. Cell Sci. 13, 637–649.

4. Meeks, J.C. and Elhai, J. (2002). Regulation of cellular differentiation in filamentous cyanobacteria in free-living and plant-associated symbiotic growth states. Microbiol. Mol. Biol. Rev. 66, 94–121.

5. Golden, J.W. and Yoon, H.S. (2003). Heterocyst development in Anabaena. Curr. Opin. Microbiol. 6, 557–563.

6. 6. Zhang, C.C., Laurent, S., Sakr, S., Peng, L., and Bédu, S. (2006). Heterocyst differentiation and pattern formation in cyanobacteria: a chorus of signals. Mol. Microbiol. 59, 367–375.

7. Kumar, K., Mella-Herrera, R.A., and Golden, J.W. (2009). Cyanobacterial Heterocysts. Cold Spring Harbor Perspectives in Biology 2, a000315–334.

8. Ishihara, JI., Tachikawa, M., Iwasaki, H., and Mochizuki, A. (2015). Mathematical study of pattern formation accompanied by heterocyst differentiation in multicellular cyanobacterium. J. Theor. Biol. 371, 9–23.

9. Asai, H., Iwamori, K., Kawai, S., Ehira, S., Ishihara, J., Aihara, K., Shouji, S., and Iwasaki, H. (2009). Cyanobacterial cell lineage analysis of the spatiotemporal hetR expression profile during heterocyst pattern formation in Anabaena sp. PCC 7120. PLoS ONE 4(10), e7371.

10. Toyoshima, M., Sasaki, N., Fujiwara, M., Ehira, S., and Ohomri, M. (2009). Early candidacy for differentiation into heterocysts in the filamentous cyanobacterium Anabaena sp. PCC 7120. Arch. Microbiol. 192, 23–31.

11. Ferimazova, N., Felcmanova, K., Setlikova, E., Kupper, H., Maldener, G., Hauska, G., Sediva, B., and Prasil, O. (2013). Regulation of photosynthesis during heterocyst differentiation in Anabaena sp strain PCC 7120 investigated in vivo at single-cell level by chlorophyll fluorescence kinetic microscopy. Photosynth. Res. 116, 79–91.

12. Kumazaki, S., Akari, M., and Hasegawa, M. (2013). Transformation of thylakoid membranes during differentiation from vegetative cell into heterocyst visualized by microscopic spectral imaging. Plant Physiol. 161, 1321–1333.

13. Sugiura, K., and Itoh, S. (2012). Single-cell confocal spectrometry of a filamentous cyanobacterium Nostoc at room and cryogenic temperature. Diversity and differentiation of pigment systems in 311 cells. Plant Cell Physiol. 53, 1492–1506.

14. Ishihara, J., Tachikawa, M., Mochizuki, A., Sako, Y., Iwasaki, H., and Morita, S. (2013). Raman imaging of the diverse states of the filamentous cyanobacteria. Nano-Bio Sensing, Imaging and Spectroscopy 8879, 88790V1–V4.

15. Dementjev, A., and Kostkeviciene, J. (2013). Applying the method of coherent anti-stokes Raman microcopy for imaging of carotenoids in microalgae and cyanobacteria. J. Raman Specrosc. 44, 973–979.

16. Tamamizu, K., and Kumazaki, S. (2019). Spectral microscopic imaging of heterocysts and vegetative cells in two filamentous cyanobacteria based on spontaneous Raman scattering and photoluminescence by 976 nm excitation. Biochim Biophys Acta Bioenerg. 1860, 78–88.

17. Huang, Y.S., Karashima, T., Yamamoto, M., and Hamaguchi, H. (2005). Molecular-level investigation of the structure, transformation, and bioactivity of single living fission yeast cells by time and space-resolved Raman spectroscopy. Biochemistry 44, 10009–10019.

18. Takanezawa, S., Morita, S., Ozaki, Y., and Sako, Y. (2015). Raman spectral dynamics of single cells in the early stages of growth factor stimulation. Biophys. J. 108, 2148–57.

19. Boucher, L.J., and Katz, J.J. (1967). The infared spectra of metalloporphyrins. J. Am. Chem. Soc. 89, 1340–1345.

20. Ogoshi, H., Saito, Y., and Nakamoto, K. (1972). Infrared spectra and normal coordinate analysis of metalloporphyrins. J. Chemical Physics 57, 4194–4202.

21. Lutz, M. (1974). Resonance Raman spectra of chlorophyll in solution. J. Raman Specrosc. 2, 497–516.

22. Koyama, Y., Umemoto, Y., Akamatsu, A., Uehara, K., and Tanaka, M. (1986). Raman spectra of chlorophyll forms. J. Mol. Struct. 146, 273–287.

23. Sashima, T., Abe, M., Kurano, H., Miyachi, S., and Koyama, Y. (1998). Changes in the carbon-carbon and carbon-nitrogen bond orders in the macrocycle of chlorophyll a upon singlet and triplet excitation as probed by resonance Raman spectroscopy of natural-abundance and singly and doubly labeled species with 15N, 13C, and 2H isotopes. J. Phys. Chem. B. 102, 6903–6914.

24. Maquelin, K., Kirschner, C., Choo-Smith, L.P., van den Braak, N., Endtz, HP., Naumann, D., and Puppels, G.J. (2002). Identification of medically relevant microorganisms by vibrational spectroscopy. J. Microbiol. Methods 51, 255–71.

25. Wood, B.R., Heraud, P., Stojkovic, S., Morrison, D., Beardall, J., and McNaughton, D. (2005). A portable Raman acoustic levitation spectroscopic system for the identification and environmental monitoring of algal cells. Anal. Chem. 77, 4955−4611.

26. Nagae, H., Kuki, M., Zhang, J.P., Sashima, T., Mukai, Y., and Koyama, Y. (2000). Vibronic coupling through the in-phase, C=C stretching mode plays a major role in the 2Ag-to 1Ag-internal conversion of all-trans-β-carotene. J. Phys. Chem. A 104, 4155−4166.

27. Kaczor, A., and Baranska, M. (2011). Structural changes of carotenoid astaxanthin in a single algal cell monitored in situ by Raman spectroscopy. Anal. Chem. 83, 7763−70.

28. Szalontai, B., Gombos, Z., and Csizmadia, V. (1985). Resonance Raman spectra of phycocyanin, allophycocyanin and phycobilisomes from blue-green alga Anacystis nidulans. Biochemical and Biophysical Research Communications 130, 358−363.

29. Szalontai, B., Gombos, Z., Csizmadia, V., and Lutz, M. (1987). The chromophore structure and chromophore-protein interactions in C-phycocyanin as studied by resonance Raman spectroscopy. Biochemica et Biophysica Acta - Bioenegetics 893, 296−304.

30. Debreczeny, M., Gombos, Z., and Szalontai, B. (1992). Surface-enhanced resonance Raman spectroscopy of phycocyanin and allophycocyanin. Eur. Biophys. J. 21, 193–198.

31. Vítek, P., Edwards, H.G., Jehlicka, J., Ascaso, C., De los Ríos, A., Valea, S., Jorge-Villar, S.E., Davila, A.F., and Wierzchos, J. (2010). Microbial colonization of halite from the hyper-arid Atacama Desert studied by Raman spectroscopy. Philos. Trans. R. Soc. A 368, 3205–3221.

32. Watanabe, M., Semchonok, D.A., Webber-Birungi, M.T., Ehira, S., Kondo, K., Narikawa, R., Ohmori, M., Boekema, E.J., and Ikeuchi, M. (2014). Attachment of phycobilisomes in an antenna-photosystem I supercomplex of cyanobacteria. Proc. Natl. Acad. Sci. USA 111, 2512–2517.

33. Peterson, R.B., Dolan, E., Calvert, H.E., and Ke, B. (1981). Energy transfer from phycobiliproteins to photosystem I in vegetative cells and heterocysts of Anabena variabilis. Biochem. Biophys. Acta. 634, 237–248.

